# PI(4,5)P2 controls slit diaphragm formation and endocytosis in *Drosophila* nephrocytes

**DOI:** 10.1101/2021.06.22.449474

**Authors:** Maximilian Gass, Sarah Borkowsky, Marie-Luise Lotz, Rita Schröter, Pavel Nedvetsky, Stefan Luschnig, Astrid Rohlmann, Markus Missler, Michael P. Krahn

## Abstract

*Drosophila* nephrocytes are an emerging model system for mammalian podocytes and podocyte-associated diseases. Like podocytes, nephrocytes exhibit characteristics of epithelial cells, but the role of phospholipids in polarization of these cells is yet unclear. In epithelia phosphatidylinositol(4,5)bisphosphate (PI(4,5)P2) and phosphatidylinositol(3,4,5)-trisphosphate (PI(3,4,5)P3) are asymmetrically distributed in the plasma membrane and determine apical-basal polarity. Here we demonstrate that both phospholipids are present in the plasma membrane of nephrocytes, but only PI(4,5)P2 accumulates at slit diaphragms. Knockdown of Skittles, a phosphatidylinositol(4)phosphate 5-kinase, which produces PI(4,5)P2, abolished slit diaphragm formation and led to strongly reduced endocytosis. Notably, reduction in PI(3,4,5)P3 by overexpression of PTEN or expression of a dominant-negative phosphatidylinositol-3-Kinase did not affect nephrocyte function, whereas enhanced formation of PI(3,4,5)P3 by constitutively active phosphatidylinositol-3-Kinase resulted in strong slit diaphragm and endocytosis defects by ectopic activation of the Akt/mTOR pathway. Thus, PI(4,5)P2 but not PI(3,4,5)P3 is essential for slit diaphragm formation and nephrocyte function. However, PI(3,4,5)P3 has to be tightly controlled to ensure nephrocyte development.

## INTRODUCTION

In *Drosophila*, pericardial nephrocytes located along the heart tube and garland nephrocytes surrounding the proventriculus filtrate the hemolymph and endocytose proteins and toxins to store the latter permanently in order to inactivate them (Weavers et al., 2009). Nephrocytes were shown to share several key features with podocytes in vertebrates, qualifying them as a model system to study mammalian podocyte function and podocyte-associated diseases (Na and Cagan, 2013; Helmstadter et al., 2017; Odenthal and Brinkkoetter, 2019). Like in podocytes, homologues of Nephrin- and Neph1 (Sticks and stones (Sns)/Hybris and Kind of irre (Kirre)/Dumbfounded) form the slit diaphragm, thereby separating the lacunae from the body cavity with hemolymph. These lacunae are formed by invaginations of the plasma membrane and form channel-like structures with both ends connected to the extra cellular space (Hochapfel et al., 2018). Due to the high endocytosis capacity in these lacunae and the expression of endocytosis receptors like Cubilin, Megalin and Amnionless, nephrocytes are used as a model system for proximal tubules of the kidney, too (Zhang et al., 2013a).

Apart from the core components of the Nephrin/Neph1 family, the slit diaphragm is stabilized by adapter proteins, e.g. the Podocin homologue Mec2 (Weavers et al., 2009; Zhang et al., 2013b) and the ZO-1 homologue Polychaetoid (Weavers et al., 2009; Carrasco-Rando et al., 2019). Furthermore, we recently showed, that regulators of classical apical-basal polarity in epithelia are partly localized to slit diaphragm complexes (Heiden et al., 2021). Knockdown studies revealed that apical polarity regulators, such as Crumbs/Stardust and the PAR/aPKC complex as well as the basolateral polarity determinants Scribble/Lethal (2) giant larvae and PAR-1 are essential for slit diaphragm formation and - at least some of them - for endocytosis (Hochapfel et al., 2017; Weide et al., 2017; Heiden et al., 2021).

In classical epithelia, these polarity regulators are targeted to either the apical (Crumbs- and PAR/aPKC-complex) or the basolateral (Scribble/Dlg/Lgl-complex, PAR-1/LKB1) plasma membrane and are essential for the establishment and maintenance of apical-basal polarity and cell-cell contacts (Flores-Benitez and Knust, 2016). However, not only proteins are involved in this process, but also distinct phospholipids are enriched either in the apical or the basolateral plasma membrane: In particular, phosphoinositol(4,5)bisphosphate (PI(4,5)P2) accumulates in the apical membrane, whereas phosphoinositol(3,4,5)trisphosphate (PI(3,4,5)P3) is preferentially found in the basolateral membrane domain (Gassama-Diagne et al., 2006; Martin-Belmonte et al., 2007). Notably, PTEN, which dephosphorylates PI(3,4,5)P3 to generate PI(4,5)P2, is recruited to the plasma membrane by PAR-3, the core scaffolding protein of the PAR/aPKC-complex (von Stein et al., 2005; Pinal et al., 2006; Feng et al., 2008). Thereby, junctionally localized PAR-3/PTEN establishes a segregation point for PI(3,4,5)P3 and PI(4,5)P2 (Martin-Belmonte et al., 2007). In turn, PAR-3 directly binds to PI(4,5)P2 and PI(3,4,5)P3, which contributes to its targeting to the plasma membrane (Krahn et al., 2010; Kullmann and Krahn, 2018a). During epithelial polarization, phosphatidylinositol-3-kinase (PI3K), which phosphorylates PI(4,5)P2 to PI(3,4,5)P3, seems to function as one of the first cues to determine the basolateral, PI(3,4,5)P3-enriched plasma membrane domain (Peng et al., 2015). Moreover, disruption of the PI(4,5)P2/PI(3,4,5)P3 balance results in severe polarity defects, suggesting a role of phospholipids as regulators of apical-basal cell polarity (Gassama-Diagne et al., 2006; Martin-Belmonte et al., 2007).

Although podocytes and nephrocytes share key features with classical epithelial cells, like cell-cell junctions and apical-basal polarity, little is known about the distribution and function of PI(4,5)P2 and PI(3,4,5)P3 in these cell types. Moreover, several studies suggest different functions of PI3K and PTEN in cultured podocytes (Huber et al., 2003a; Bridgewater et al., 2005; Zhu et al., 2008; Wang et al., 2014), but the role of these key enzymes *in vivo* is still unclear. Therefore, the aim of this study was to investigate the subcellular accumulation of these two phospholipids as well as their function in slit diaphragm assembly and nephrocyte development.

## RESULTS

### PI(4,5)P2 but not PI(3,4,5)P3 is enriched at slit diaphragms

In classical epithelia, PI(4,5)P2 is enriched in the apical plasma membrane, whereas PI(3,4,5)P3 accumulates in the basolateral plasma membrane (Martin-Belmonte et al., 2007). In contrast, nothing is known about the distribution of specific phospholipids in mammalian podocytes or *Drosophila* nephrocytes. Therefore, we first investigated the distribution of PI(4,5)P2 and PI(3,4,5)P3 in *Drosophila* garland nephrocytes by expressing fusion proteins consisting of a fluorescent protein and a Pleckstrin homology (PH) domain, which preferentially bind to PI(4,5)P2 (PH domain of PLCδ (Verstreken et al., 2009)) or to PI(3,4,5)P3 (PH domain of Akt1, this study).

mCherry-PH(PLCδ) is substantially associated with the plasma membrane (Fig. 1A-B) but it is also found in intracellular pools, partly associated with vesicular structures. Surface views reveal that its cortical association form strand-like structures, which to some extent co-stain with endogenous Sns, a marker for slit diaphragms (Fig. 1B). In contrast, PH(Akt)-GFP is only weakly associated with the plasma membrane but also shows a cytoplasmic and vesicular-associated distribution (Fig. 1C). Nonetheless, surface views show a strand-like pattern too, but these strands do not co-localize with Sns (Fig. 1D) but are rather found between the Sns-strands. These findings suggest, that PI(4,5)P2 in the plasma membrane accumulates at slit diaphragms, whereas PI(3,4,5)P3 is enriched in the free plasma membrane between slit diaphragms.

**Figure 1.**
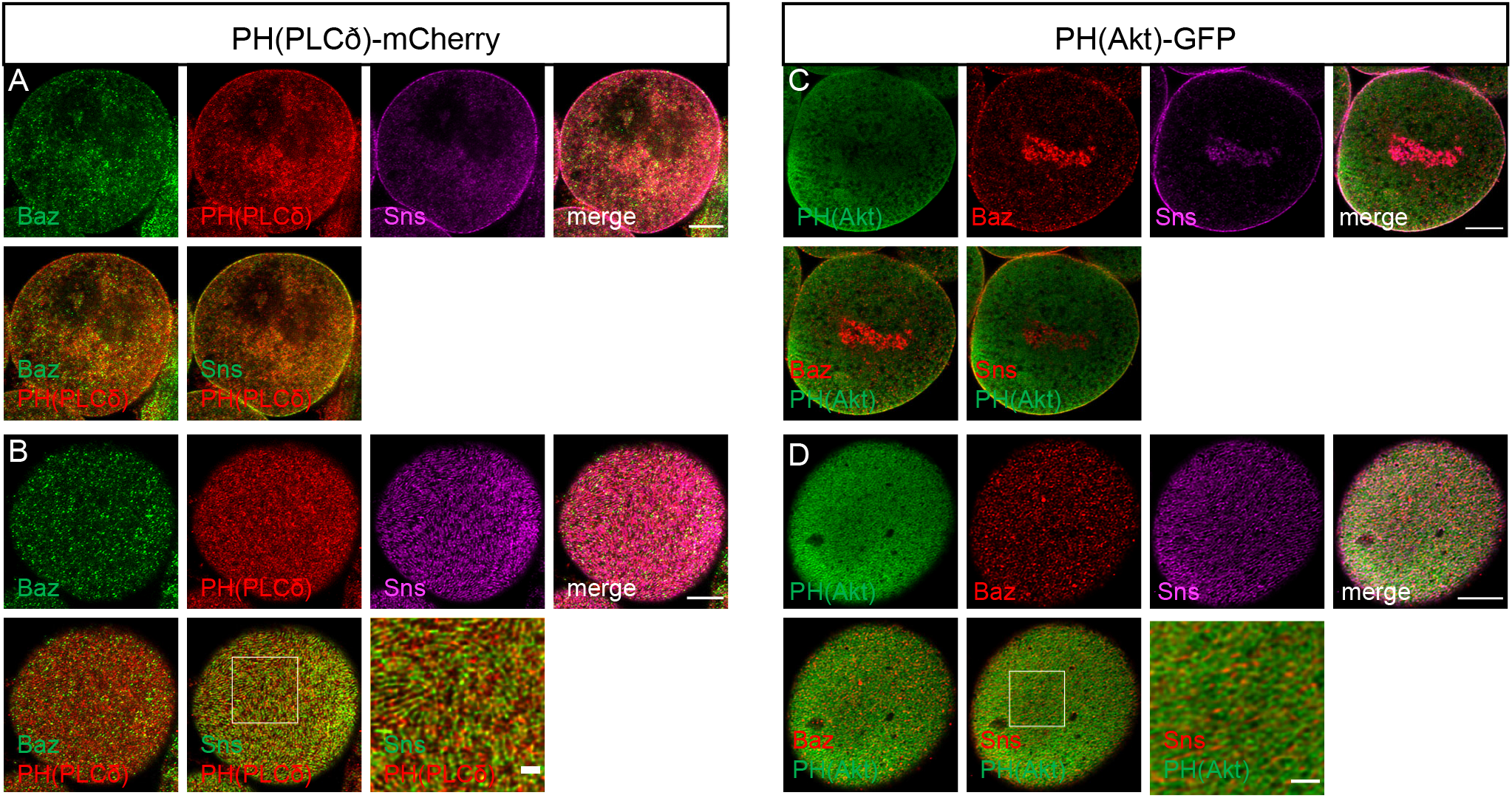
PI(4,5)P2 accumulates at slit diaphragms whereas PI(3,4,5)P3 is found at the free plasma membrane. (A-D) Garland nephrocytes expressing either UAS::PH(PLCδ)-mCherry for labelling PI(4,5,)P2 or UAS::PH(Akt)-GFP to visualize PI(3,4,5)P3 were dissected from 3^rd^ instar larvae, fixed and stained with the indicated antibodies. A and C are sections through the equatorial region of the nephrocyte and B/D are onviews onto the surface of these nephrocytes. Scale bars are 5μm and 1μm in insets.

### Impaired PI(4,5)P2 production results in strong developmental and slit diaphragm defects

In order to test whether PI(4,5)P2 is essential for nephrocyte development and function, in particular regarding slit diaphragm assembly and maintenance, we used RNA-interference (RNAi) to knockdown the ubiquitously expressed PI(4)P5-Kinase Skittles (Sktl), which is responsible for converting PI(4)P to PI(4,5)P2 using the nephrocyte-specific driver line sns::GAL4. In *Drosophila*, Sktl has been described to regulate apical-basal polarity by targeting PAR-3 to the apical junctions in follicular epithelial cells (Claret et al., 2014) and to the anterior cortex in the oocyte (Wen et al., 2020). In tracheal tubes, Sktl-produced PI(4,5)P2 was proposed to recruit the formin Diaphanous to the apical membrane (Rousso et al., 2013). In nephrocytes, Sktl partly colocalizes with Sns at slit diaphragms (Fig. S1A), opening the possibility of a local accumulation of PI(4,5)P2 in microdomains of the plasma membrane at slit diaphragms. Indeed, impaired expression of Sktl resulted in dramatic morphological changes with fused nephrocytes (Fig. 2B compared to control RNAi in 2A). Furthermore, the typical strand-like structures of Sns-labelled slit diaphragm observed at the surface of control nephrocytes was completely abolished in Sktl-RNAi expressing nephrocytes, resulting in a dispersion of Sns to intracellular puncta (Fig. 2A-D). Besides Sns, the basal polarity determinant Talin and the apical polarity regulator PAR-3 (Bazooka (Baz) in Drosophila) are lost from the cortex, too. In contrast to impaired PI(4,5)P2 levels, overexpression of Sktl in order to increase PI(4,5)P2 did not affect nephrocyte morphology or slit diaphragm assembly (Fig. S1B, quantified in Fig. 2E), although the amount of PI(4,5)P2 seemed to be significantly increased, as demonstrated by enhanced accumulation of mCherry-PH(PLCδ) at the plasma membrane (Fig. S1C).

**Figure 2.**
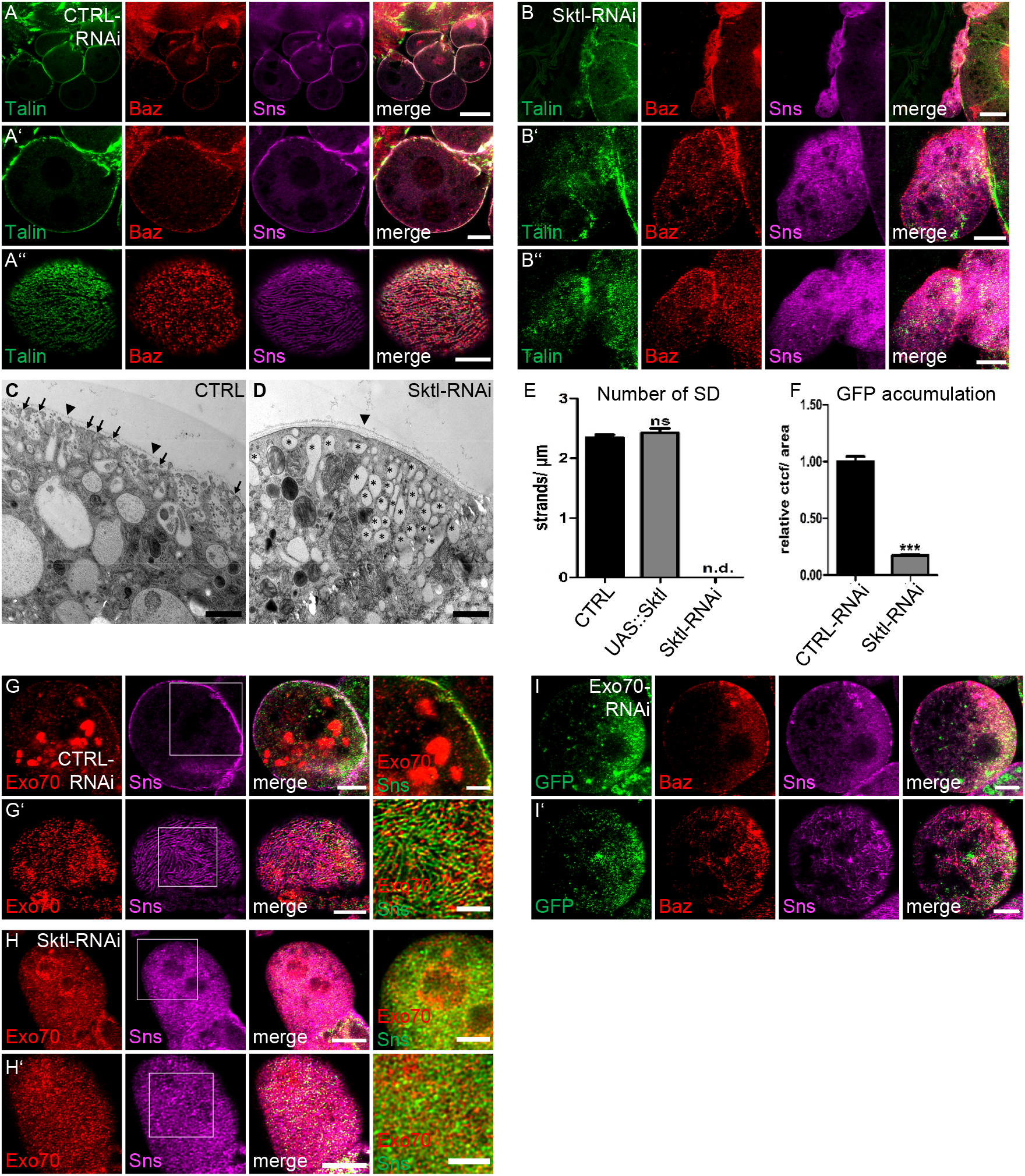
PI(4,5)P2 produced by Skittles is essential for slit diaphragm formation and endocytosis. (A-B) Garland nephrocytes from 3^rd^ instar larvae expressing either control RNAi (A) or Sktl-RNAi (B) were stained with the indicated antibodies. (C-D) Transmission electron microscopy of garland nephrocytes of control third instar larvae (C) and Sktl-RNAi-expressing larvae (D). Some slit diaphragms were labeled with arrows in control nephrocytes. Slit diaphragms were absent in Sktl-RNAi expressing nephrocytes. Arrow heads mark the basement membrane. Asterisks indicate electron-bright vesicles, which resembles lacunae in control nephrocytes. (E) Slit diaphragms of nephrocytes expressing Sktl or control were quantified from surface views. For this, a 5μm line perpendicular to the Sns-strands was drawn and the number of strands quantified. 5 lines/nephrocyte and at least 5 nephrocytes were quantified per genotype. Sktl-RNAi expressing nephrocytes were not characterized as they did not display detectable Sns strands at the surface but exhibited a rather diffuse Sns staining. (F) Endocytosis of a secreted ANP-2xGFP by garland nephrocytes expressing the indicated RNAi’s was quantified as described in the methods section. At least 100 nephrocytes from at least 15 different larvae were evaluated. (G-H) Nephrocytes expressing control RNAi (G) or Sktl RNAi (H) were co-stained with Exo70 and Sns. (I) Immunostainings of nephrocytes expressing Exo70-RNAi. Scales bars are 15μm in A and B, 5μm in A, A’’, B’, B’’ and G-I, 2.5μm in insets in G and H and 1μm in C-D. Error bars are standard error of the means. Significance was determined by Mann-Whitney test: *** p<0.001, ** p<0.01,* p<0.05. n.s. not significant.

### Skittles is essential for slit diaphragm assembly by regulating exocytosis

Analysis of Sktl-RNAi expressing nephrocytes by electron microscopy confirmed an almost complete absence of slit diaphragms (Fig. 2D compared to control in C). Notably, these nephrocytes do not form regular lacunae but accumulate large electron-light vesicles below the plasma membrane (marked with asterisks in Fig. 2D). This phenotype suggests severe defects in exocytosis, which is essential for the delivery of transmembrane proteins of the slit diaphragm complex (Sns, Kirre and Crb). During exocytosis, clustering of PI(4,5)P2 facilitates the docking of the exocyst complex to the plasma membrane by direct binding of its components Exo70 and Sec3 in yeast and in mammalian cells (He et al., 2007; Liu et al., 2007; Zhang et al., 2008; Martin, 2015). In a second step, PI(4,5)P2 is also essential for vesicle fusion and several proteins involved in regulation of fusion directly interact with PI(4,5)P2 (reviewed by Schink et al., 2016). In order to test whether Sktl-produced PI(4,5)P2 recruits Exocyst complex components in nephrocytes, we stained for endogenous Exo70. In control nephrocytes, apart from intracellular giant vesicles, a substantial pool of Exo70 was found at the plasma membrane, co-localizing with Sns (Fig. 2G). In contrast, it displayed a diffuse localization with some perinuclear accumulation in Sktl-RNAi expressing nephrocytes (Fig. 2H). Moreover, downregulation of the exocyst complex components Exo70 and Sec3 resulted in similar loss of slit diaphragms as Sktl-RNAi (Fig. 2I and Fig. S1D), which is in line with a recent study reporting a crucial role of the exocyst complex in slit diaphragm formation/maintenance (Wen et al., 2020).

### Decreased PI(4,5)P2 levels impair endocytosis in nephrocytes

Apart from exocytosis, PI(4,5)P2 also regulates clathrin-dependent and -independent endocytosis by recruiting several proteins involved in early steps of endocytosis to the plasma membrane and by inducing actin remodeling during micropinocytosis (reviewed by Schink et al., 2016). In nephrocytes, endocytosis is essential for the uptake of filtrated proteins, toxins and metabolites, which are then stored and inactivated. Disturbance of slit diaphragm formation as well as of endocytic receptors and proteins involved in the endocytosis machinery have been reported to reduce endocytosis (Soukup et al., 2009; Weavers et al., 2009; Zhuang et al., 2009; Zhang et al., 2013a; Hochapfel et al., 2017; Weide et al., 2017; Hermle et al., 2018; Kampf et al., 2019; Troha et al., 2019). In order to test, whether PI(4,5)P2 is essential for endocytosis in nephrocytes, we quantified the accumulation of secreted ANP-2xGFP (Heiden et al., 2021) which is secreted into the hemolymph, filtrated by nephrocytes and taken up by endocytosis. Indeed, downregulation of Sktl in nephrocytes, reducing PI(4,5)P2 levels, resulted in a strong decrease of ANP-2xGFP accumulation in nephrocytes, consistent with impaired endocytosis (Fig. 2F). This is in line with reports from the *Drosophila* oocyte, where Sktl is essential for Rab5-mediated endocytosis of yolk protein (Compagnon et al., 2009).

### PI(3,4,5)P3 is not essential for nephrocyte function but ectopic production results in dominant negative effects

In contrast to PI(4,5)P2, reducing PI(3,4,5)P3 by overexpression of PTEN or expression of a dominant negative version of PI3K (PI3K-DN) did not affect nephrocyte morphology or slit diaphragm formation (Fig. S2A-B and Fig. 3F). However, overexpression of a constitutively active PI3K (PI3K-CA), which is targeted to the plasma membrane by attachment of a prenylation anchor (CAAX-motif), in nephrocytes resulted in a strong fusion phenotype and a disturbed pattern of slit diaphragms (Fig. 3A-D, quantified in 3F). Notably, PI3K-CA-expressing nephrocytes are larger than control nephrocytes (Fig. 3G). In addition to slit diaphragm defects, overexpression of PI3K-CA resulted in a drastic decrease in ANP-2xGFP uptake, suggesting a defect in endocytosis (Fig. 3H).

**Figure 3.**
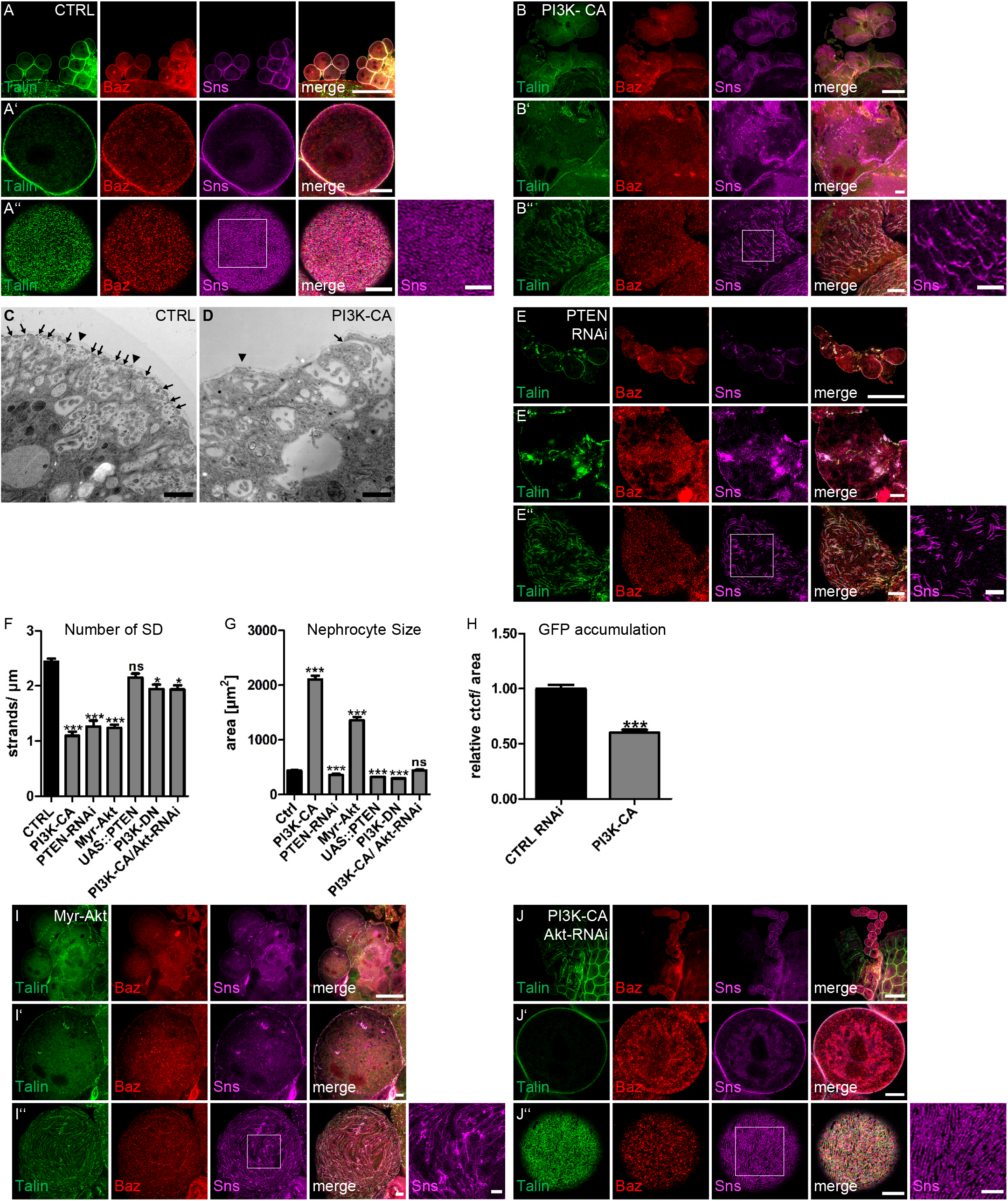
PI(3,4,5)P3 is not essential for slit diaphragm assembly and maintenance. (A-B) Garland nephrocytes from 3^rd^ instar larvae either of controls (sns::GAL4 crossed with the empty attP40 line, A) or of animals expressing a constitutively activated Pi3K (Pi3K-CA, B) in nephrocytes were stained with the indicated antibodies. (C-D) Transmission electron microscopy of garland nephrocytes of control third instar larvae (C) and PI3K-CA expressing larvae (D). Slit diaphragms were labeled with arrows and arrow heads mark the basement membrane. (E) Immunostainings of nephrocytes expressing RNAi against PTEN. (F) Slit diaphragms of nephrocytes expressing the indicated transgenes were quantified from surface views. For this, a 5μm line perpendicular to the Sns-strands was drawn and the number of strands quantified. 5 lines/nephrocyte and at least 5 nephrocytes were quantified per genotype. (G) The size of nephrocytes expressing the indicated transgenes was quantified by measuring the cell area of equatorial sections. At least 120 nephrocytes from at least 10 larvae were quantified. (H) Endocytosis of a secreted ANP-2xGFP by garland nephrocytes expressing the indicated controls was quantified as described in the methods section. At least 100 nephrocytes from at least 15 different larvae were evaluated. (I) Myr-Akt expressing nephrocytes were stained with the indicated antibodies. (J) Immunostainings of nephrocytes expressing Pi3K-CA together with RNAi targeting Akt. Scale bars are 50μm in A, B, E, I and J, 5μm in A’, A’’, B’, B’’, E’, E’’, I’, I’’, J’ and J’’, 2.5μm in insets in A’’, B’’, E’’, I’’, J’’ and 1μm in C and D. Error bars are standard error of the means. Significance was determined by Mann-Whitney test: *** p<0.001, ** p<0.01,* p<0.05. n.s. not significant.

Like PI3K-CA, enhanced accumulation of PI(3,4,5)P3 by knockdown of PTEN resulted in similar but milder phenotypes regarding slit diaphragms, whereas cell size was not increased (Fig. 3E-G). This is likely due to the limited abundance of PI(3,4,5)P3 within the plasma membrane. Ectopic production of PI(3,4,5)P3 from PI(4,5)P2 by PI3K-CA likely produces higher levels of PI(3,4,5)P3 in the plasma membrane due to the larger pool of PI(4,5)P2 (Lemmon, 2008), whereas inhibition of dephosphorylation of PI(3,4,5)P3 to PI(4,5)P2 only moderately increases PI(3,4,5)P3 levels in the plasma membrane. These data suggest that slit diaphragm assembly might be more sensitive to enhanced PI(3,4,5)P3 level than cell size regulation.

### Phenotypes of increased PI(3,4,5)P3 are accomplished by the mTOR pathway

Increased PI(3,4,5)P3 in the plasma membrane leads to activation of the Akt/mTOR signaling cascade, which, among various other functions, results in cell survival and increased cell size and proliferation (reviewed by Saxton and Sabatini, 2017). In order to test whether the phenotypes observed in nephrocytes expressing PI3K-CA are caused by ectopic Akt/mTOR activation, we introduced a constitutively active variant of Akt (Myr-Akt), which is recruited to the plasma membrane and activated independently of PI(3,4,5)P3 due to the fusion of a myristoylation-signal (Stocker et al., 2002). Indeed, these nephrocytes mimicked the PI3K-CA overexpression phenotype with disrupted slit diaphragms, increased size and fusion phenotypes (Fig. 3F, G, I). However, cell size of Myr-Akt expressing nephrocytes was not as strongly increased as in PI3K-CA expressing ones (albeit higher than in case of PTEN-RNAi), whereas slit diaphragm assembly is severely disturbed and comparable with Pi3K-CA and PTEN-RNAi-expressing nephrocytes. Thus, these data provide additional support to the notion that slit diaphragm assembly and size regulation show different susceptibility to levels of PI(3,4,5)P3. To further substantiate our hypothesis that the defects observed in PI3K-CA expressing nephrocytes are due to ectopic activation of Akt/mTOR signaling upon increased levels of PI(3,4,5)P3, we knocked down Akt or *Drosophila* Tor (dTOR) in PI3K-CA expressing nephrocytes. As depicted in Fig. 3F,G,J and Fig. S2C, downregulation of Akt or dTOR rescued to a large extent the slit diaphragm defects as well as size differences in PI3K-CA expressing nephrocytes, confirming that the dominant negative function of PI(3,4,5)P3 is mediated by the Akt/mTOR-pathway.

### Changes in PI(4,5)P2 and PI(3,4,5)P3 levels cause rapid defects

In order to elucidate whether slit diaphragm defects are established early in development during formation of nephrocytes or whether PI(4,5)P2 and PI(3,4,5)P3 levels are also essential for the turnover and maintenance of slit diaphragms, we used a temperature-sensitive GAL80 (GAL80ts), which suppresses GAL4 activity at the permissive temperature at 18°C. After molting to L3, larvae were shifted to 29°C for 24h prior to dissection, inactivating the GAL80 and thus releasing GAL4, which induces the UAS-transgene. In Sktl-RNAi expressing nephrocytes dissected from animals raised under these conditions, we observed similar defects in morphology as well as impaired Sns strands (Fig. 4A-C), indicating that PI(4,5)P2 is essential for the turnover/maintenance after the initial establishment of slit diaphragms during the development of nephrocytes. In contrast, short-term induction of Pi3K-CA did not produce phenotypes comparable to continuous expression of this transgene (Fig. 4D-F), indicating that the Akt/mTOR-mediated effect of ectopic PI(3,4,5)P3 production is either critical during nephrocyte development or it takes longer time to get established, presumably due to the delay upon transcriptional reprogramming of the cell as a consequence of mTOR target activation.

**Figure 4.**
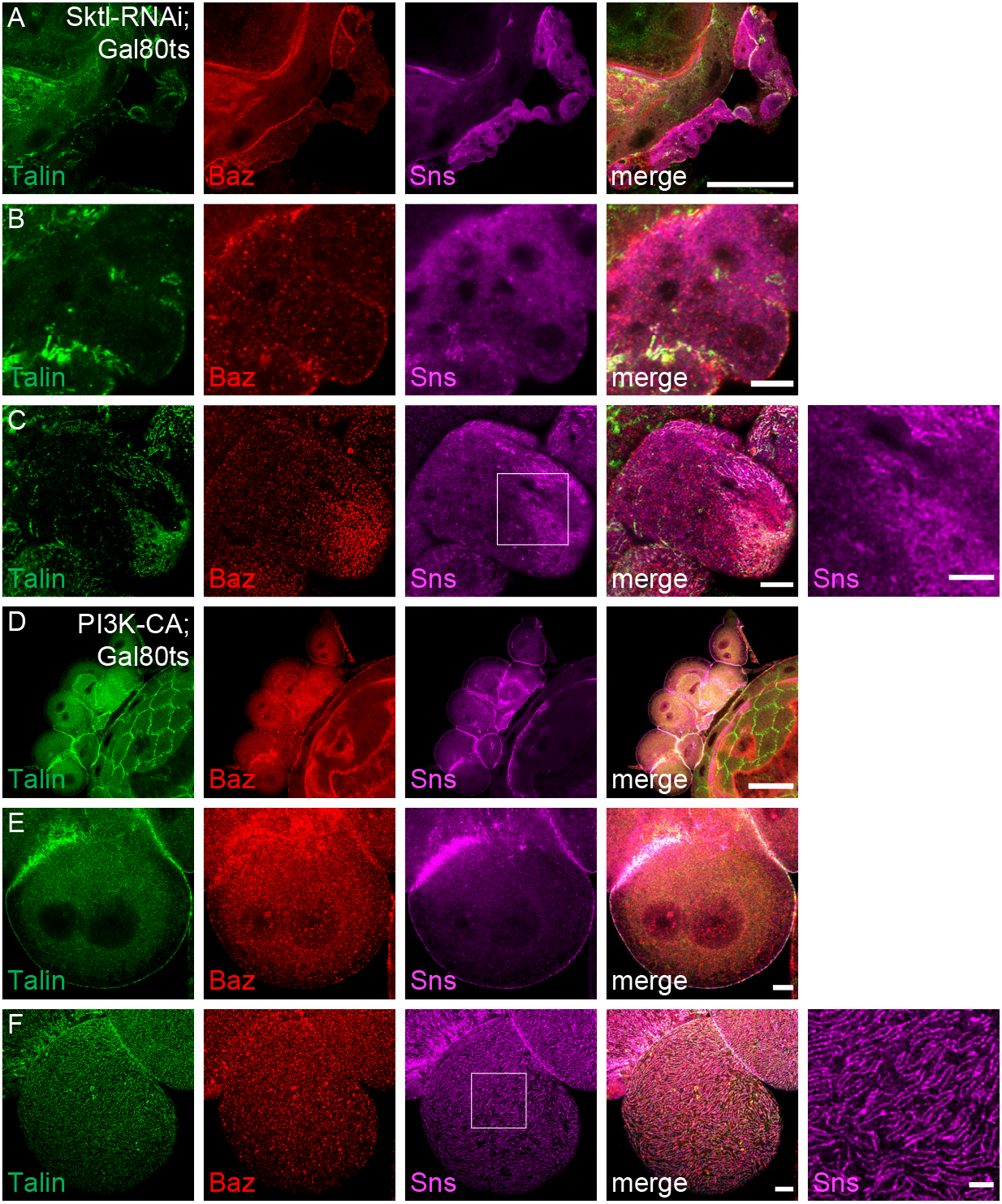
Rapid effects of PI(4,5)P2 reduction and PI(3,4,5)P3 accumulation. (A-F) Immunostainings of nephrocytes from 3^rd^ instar larvae expressing GAL80ts together with sns::GAL4 and Sktl-RNAi (A-C) or Pi3K-CA (D-F), which were first raised at 18°C in order to suppress expression of the transgenes. In L3, larvae were shifted to 29°C for 24h prior to dissection in order to induce expression of the transgenes. Scale bars are 50μm in A and D, 5μm in B,C, E and F and 2.5μm in insets in C and F.

## DISCUSSION

Our findings demonstrate that PI(4,5)P2, but not PI(3,4,5)P3 is essential for nephrocyte function and slit diaphragm formation. Of note, PI(4,5)P2 is not evenly distributed in the entire plasma membrane but displays a strand-like pattern, partly colocalizing with Sns as a maker for slit diaphragms. Although PI(4,5)P2 has been found in other cell types at the entire plasma membrane – or, in epithelial cells, enriched in the apical plasma membrane domain - there is increasing evidence that this phospholipid is concentrated in distinct microdomains of the plasma membrane (discussed by Kwiatkowska, 2010; Kolay et al., 2016): In cultured fibroblasts, freeze-fracture membrane preparation and subsequent electron microscopy revealed three distinct pools of PH(PLCδ) at the rim of caveolae, in coated pits and at the free plasma membrane (Fujita et al., 2009). Notably, these three pools exhibited different kinetics upon regulatory stimuli, suggesting different types of regulation. PI(4,5)P2 was also reported to accumulate in lipid rafts of distinct (phospho)lipid and cholesterol composition within the plasma membrane, promoting local actin remodeling or receptor clustering (Golub and Caroni, 2005; Szymanska et al., 2009; Capuano et al., 2012). Sarmento et al. observed a Ca(2+)-dependent PI(4,5)P2 clustering in liposomes *in vitro* under physiological Ca(2+) and PI(4,5)P2 concentrations (Sarmento et al., 2014). Thus, PI(4,5)P2 may accumulate in distinct microdomains of the plasma membrane adjacent to slit diaphragms in order to regulate vesicle trafficking – to the plasma membrane by inducing fusion of vesicles and from the plasma membrane by regulating endocytosis. The dramatic phenotypes observed in Sktl-RNAi expressing nephrocytes underline the critical role of PI(4,5)P2 as an important regulator in these processes. Notably, the human homologue of Sktl, PIP5Kα, was described to be recruited by the Chloride Intracellular Channel 5 (CLIC5A) to cortical Ezrin, inducing clusters of PI(4,5)P2 in the plasma membrane of COS-7 cells (Al-Momany et al., 2014). In podocytes, Ezrin is part of the Ezrin-NHERF2-Podocalyxin complex, an essential component of the glycocalyx. Furthermore, in glomeruli of CLIC5A-deficient mice, cortical Ezrin/NHERF2 as well as glomerular Podocalyxin are reduced (Al-Momany et al., 2014). Another hint to an important role of PI(4,5)P2 in regulating podocyte morphology comes from a study reporting that the PI5P-Phosphatase Ship2 can be recruited and activated by Nephrin via Nck-Pak1-Filamin in cultured human podocytes (Venkatareddy et al., 2011). Ship2 dephosphorylates PI(3,4,5)P3 to PI(3,4)P2, thus its activation by Nephrin in this systems results in an increase of PI(3,4)P2, which activates Lamellipodin, a regulator of Ena/Vasp proteins, resulting in the formation of lamellipodia. Finally, the Nephrin/Ship2 interaction was increased in a podocyte injury model *in vivo*, suggesting that lamellipodia formation upon Nephrin-mediated Ship2-activation contributes to foot process effacement observed upon podocyte damage. However, it remains unclear how the Ship2-regulated balance between PI(3,4,5)P3 and PI(3,4)P2 at the Nephrin-complex contributes to slit diaphragm assembly/maintenance and podocyte function under physiological conditions.

PI(4,5)P2 as well as PI(3,4,5)P3 are capable of regulating the actin cytoskeleton by recruiting and activating the small GTPases Rac1 and Cdc42 as well as proteins of the WASP family (Prehoda et al., 2000; Rohatgi et al., 2000; Padrick and Rosen, 2010). Notably, a coordinated actin cytoskeleton remodeling is essential for cortical Nephrin localization and slit diaphragm assembly in *Drosophila* nephrocytes (Muraleedharan et al., 2018; Bayraktar et al., 2020) as well as in mammalian podocytes (Perico et al., 2016). *Vice versa*, activated Nephrin recruits PI3K resulting in Rac1 activation, actin branching and lamellipodia formation in cultured rat podocytes (Zhu et al., 2008). Notably, PTEN is downregulated in podocytes of patients suffering from diabetic nephropathy and inhibition or podocyte-specific knockout of PTEN in mice results in cytoskeleton rearrangements, foot process effacement and proteinuria (Lin et al., 2015).

Apart from their impact on the actin cytoskeleton, PI(4,5)P2 and PI(3,4,5)P3-activated Rac1/Cdc42 and actin regulators are essential for remodeling and stability of tight junctions as well as adherens junctions in classical epithelia (reviewed by Shewan et al., 2011). Increasing evidence suggests that the slit diaphragms connecting the foot processes of neighboring podocytes emerge from transformation of the tight junctions of the epithelial podocyte progenitor cells (Simons et al., 2009). Indeed, several proteins of the adherens- and tight junctions can also be found to be components of the slit diaphragm, e.g. ZO-1, Crumbs, PAR/aPKC-complex (Huber et al., 2003b; Hartleben et al., 2008; Hirose et al., 2009; Weavers et al., 2009; Hartleben et al., 2013; Itoh et al., 2014; Satoh et al., 2014; Hochapfel et al., 2017; Weide et al., 2017; Carrasco-Rando et al., 2019; Moller-Kerutt et al., 2021). Thus, it is likely that changes in PI(4,5)P2 and PI(3,4,5)P3 affect slit diaphragm formation and maintenance/stability like they affect adherens junctions/tight junctions in classical epithelia.

## MATERIALS AND METHODS

### *Drosophila* stocks and genetics

Fly stocks were cultured on standard cornmeal agar food and maintained at 25°C. For downregulation or overexpression of specific genes for immunostainings and electron microscopy, *sns::GAL4* (Zhuang et al., 2009), was crossed with the following lines: UAS::Akt-RNAi (#103703), UAS::Exo70-RNAi (#103717), UAS::Or83b-RNAi (negative control, #100825), UAS::PTEN-RNAi (#01475), UAS::Sec3-RNAi (#108085), UAS::Sktl-RNAi (#101624) (provided by Vienna *Drosophila* Resource Center, Austria), UAS::PI3K92E-CAAX (PI3K-CA, #8294), UAS::PI3K92E.A2860C (PI3K-DN, #8289), UAS:Sktl (#39675), UAS::PH(PLCδ)-mCherry (#51658), tubP::GAL80ts (65406), UAS::dTOR-RNAi (#34639) (all obtained from Bloomington stock center). UAS::Myr::Akt was provided by Hugo Stocker (Stocker et al., 2002) and UAS::Myc-Sktl was obtained from Sandra Claret (Jouette et al., 2019). UASt::PTEN was established by PhiC31-Integrase Insertion using attP86F. UAS::PH(Akt)-GFP was constructed by fusing the PH domain of mammalian Akt1 to the N-terminus of GFP in the pUASt-vector. Transgenic flies were generated by P-element-mediated germ line transformation. An insertion on second chromosome was used in this study. For all RNAi and overexpression experiments, crosses were kept for 3 days at 25°C and larvae subsequently shifted to 29°C, in order to obtain maximum expression. PH(PLCδ)-mCherry was expressed at 25°C, PH(Akt)-GFP was analyzed at 18°C, 21°C and 25°C, with best results at 18°C, because at higher temperature, the expression of the chimeric protein was too strong and found overall the cell, likely due to the limited amount of PI(3,4,5)P3 to bind to.

### Endocytosis assays

For the ANP-2xGFP accumulation assay, garland nephrocytes from wandering third instar larvae were dissected in HL3.1 saline (Feng et al., 2004), fixed in 4% PFA in PBS for 10min, stained with DAPI for 20min, washed with PBS, and mounted in Mowiol. ANP-2xGFP accumulation per nephrocyte area (CTCF = Corrected Total Cell Fluorescence) was analyzed and quantified with ImageJ after subtracting the autofluorescent background of dissected larvae. For each genotype, at least 100 nephrocytes of 15 independent larvae were quantified.

### Immunohistochemistry

Garland nephrocytes were dissected as described above and heat-fixed for 20 seconds in boiling heat fix saline (0.03% Triton-X100). Subsequently, nephrocytes were washed three times in PBS + 0.2% Triton X-100 and blocked with 1% BSA for 1h, incubated over night with primary antibodies in PBS + 0.2% Triton X-100 + 1% BSA, washed three times and incubated for 2h with secondary antibodies. After three washing steps and DAPI-staining, nephrocytes were mounted with Mowiol. Primary antibodies used were as follows: anti Baz (1:250, Kullmann and Krahn, 2018b), rabbit anti Exo70 (1:500, Koon et al., 2018), goat anti GFP (1:500, #600-101-215, Rockland), mouse anti Myc (1:100, 9E10, Developmental Studies Hybridoma Bank (DSHB)), chicken anti Sns (1:1000, Hochapfel et al., 2017), mouse anti Talin (1:20, E16B, DSHB). Secondary antibodies conjugated with Alexa 488, Alexa 568 and Alexa 647 (Life technologies) were used at 1:400. Images were taken on a Leica SP8 confocal microscope using lightning program and processed using ImageJ.

### Transmission electron microscopy

Garland nephrocytes of third instar larvae were dissected in HL3.1 saline, high pressure frozen (EM-PACT2, Leica, Wetzlar, Germany), freeze-substituted in acetone / 1% OsO4 / 5% H_2_O / 0.25% uranyl acetate (AFS2, Leica, Wetzlar, Germany) and embedded in Epon. For transmission electron microscopy, 70nm thick sections were cut using an ultramicrotome (Leica UC7, Wetzlar, Germany). All samples were imaged with a transmission electron microscope (ZEISS, Libra 120, Germany).

## Acknowledgements

We thank E. Chan, S. Claret, the Bloomington *Drosophila* stock center at the University of Indiana (USA), the Vienna Drosophila Resource Center (Austria) and the Developmental Studies Hybridoma Bank at the University of Iowa (USA) for providing reagents. We also thank Kerstin Seiling for technical assistance with electron microscopic work. This work was supported by grants of the German research foundation (DFG) to M. P. K. (CRC1348-A05), M.M. (CRC1348-A03) and S.L. (CRC1348-B10).

## Author contributions

M.G., S.B. M.-L.L. performed the experiments and analyzed the data, S.L. established the UAS::Akt-GFP line and revised the manuscript, A.R., R.S. and M.M. performed electron microscopy analysis and revised the paper, P.N. and M.P.K. supervised the experiments and wrote the manuscript.

## Competing interests

The authors declare no competing interests.

---

**Figure S1.**
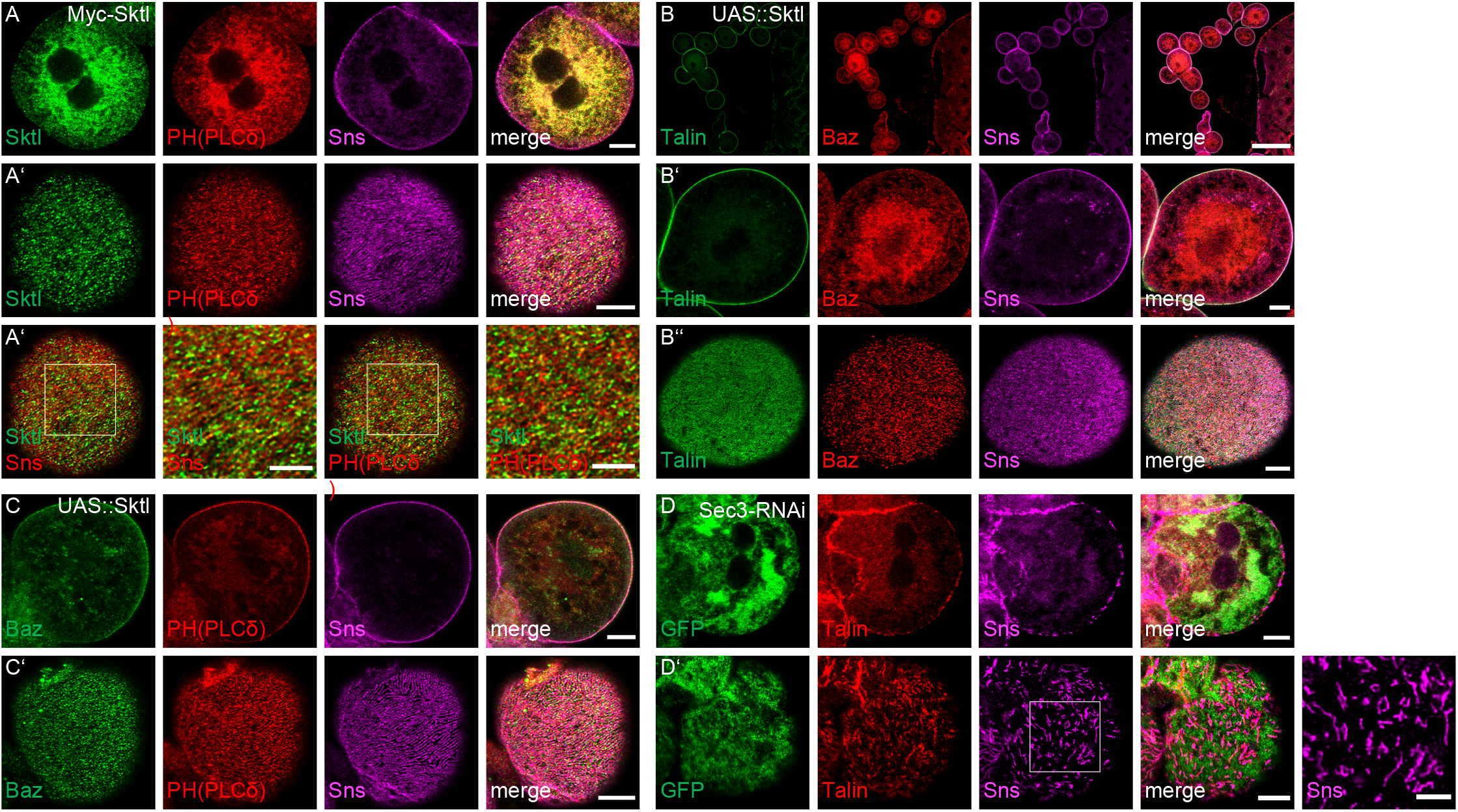
Overexpression of Sktl does not affect slit diaphragm assembly. Related to Fig. 2. (A) Immunostaining of Skittles-GFP, Sns and Baz in nephrocytes. (B) Nephrocytes overexpressing Sktl were stained with the indicated antibodies. (C) Overexpression of Sktl results in increased accumulation of PH(PLCδ)-mCherry at the plasma membrane. Scale bars are 5μm in A, A’’, B’, B’’, C, C’, 50μm in B and 2,5μm in insets in A’.

**Figure S2.**
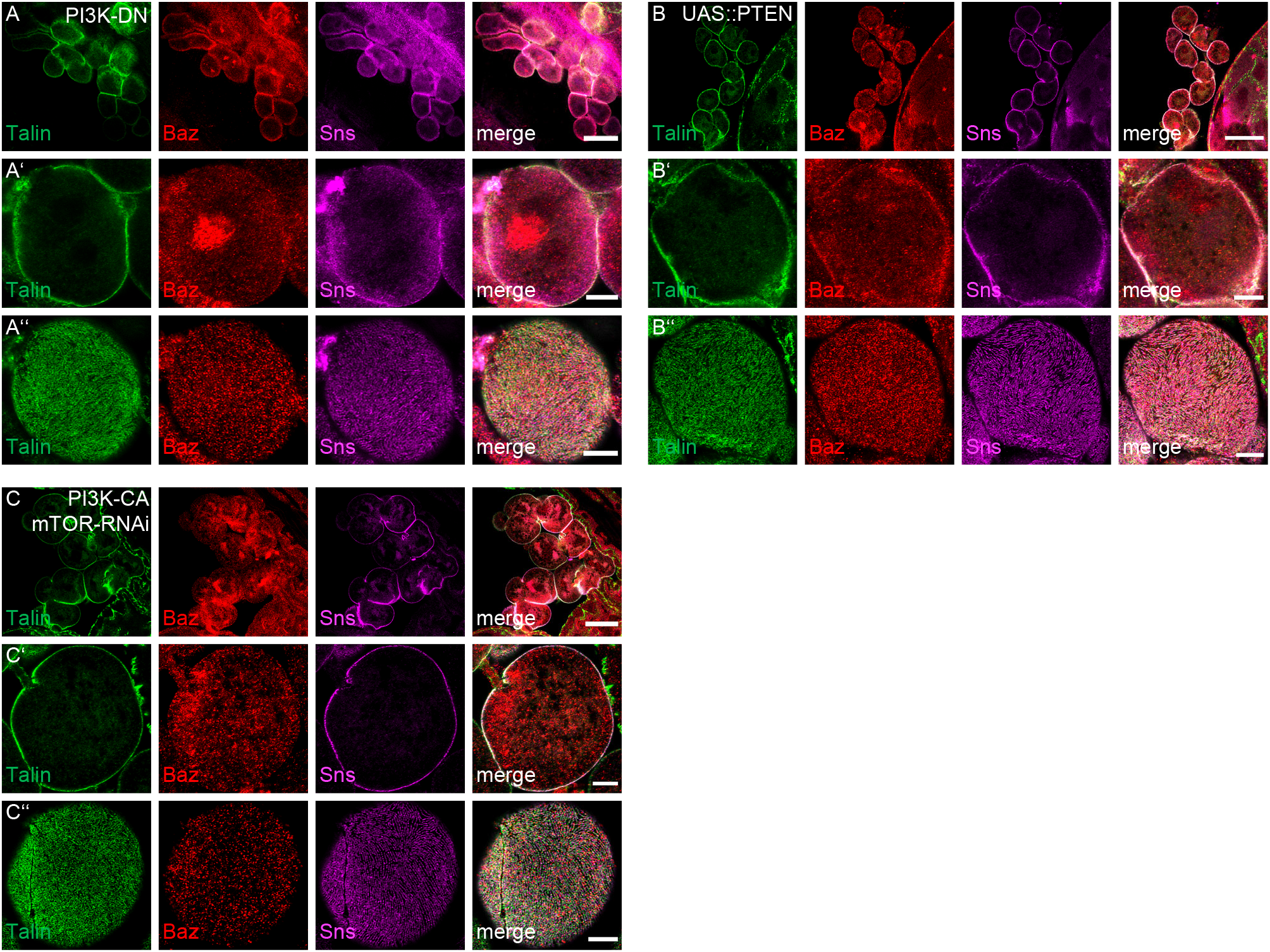
Decrease of PI(3,4,5)P3 does not affect slit diaphragms. Related to Fig. 3. (A-B) Nephrocytes overexpressing a dominant negative Pi3K (Pi3K-DN, A) or PTEN (B) were stained with the indicated antibodies. (D) RNAi targeting dTOR was expressed in nephrocytes with expression of Pi3K-CA. Scale bars are 25μm in A, B and C and 5μm in A’, A’’, B’, B’’, C’ and C’’.

## Notes

### Competing Interest Statement

The authors have declared no competing interest.

